# Gelatinous matrix, an original strategy to cope with oligotrophy in Nassellaria (Radiolaria)

**DOI:** 10.1101/2024.01.22.576765

**Authors:** Natalia Llopis Monferrer, Sarah Romac, Manon Laget, Yasuhide Nakamura, Tristan Biard, Miguel M. Sandin

## Abstract

Radiolaria are heterotrophic protists abundant in the world’s oceans playing important roles in biogeochemical cycles. Some species host photosynthetic algae also contributing to primary production. Such mixotrophic behaviour is believed to explain their ecological success in oligotrophic waters, notably Collodaria, exclusively mixotrophic radiolarians within a gelatinous matrix. Yet, our understanding of Radiolaria ecology is limited to direct observations, as they have so far withstood reproduction in culture and their genomes are unexplored. Sampling oligotrophic California Current communities revealed an abundant, rarely observed population of Nassellaria of the genus *Phlebarachnium*, characterized to live within a gelatinous matrix along with other Radiolaria. Phylogenetic reconstruction of the ribosomal DNA suggests that these distantly related lineages within Nassellaria independently developed the ability to produce a gelatinous matrix ∼150 million years ago. By matching physical samples with their genetic signature, we identified these rarely observed organisms in global metabarcoding datasets, revealing strong biogeographic affinity to oligotrophic water masses. Global ocean co-occurrence networks showed that Radiolaria with a gelatinous matrix have a distinct biogeography compared to those without the matrix. Results suggest that the gelatinous matrix is an adaptation to oligotrophic waters, but further research is needed to evaluate similarities between the gelatinous matrices across different Radiolaria groups. This strategy could increase the effective volume to weight ratio favoring prey capture and create a favorable microenvironment for symbionts, enhancing ecological success in nutrient-depleted waters. This study advances our understanding of eukaryotic diversity evolution, emphasizing specific advantages of certain adaptations, specifically when evolution occurs independently across lineages.

## Introduction

Among marine planktonic eukaryotic organisms, Radiolaria are diverse and highly abundant protists, widely distributed in the ocean, from tropical to polar waters [1]. These protists exhibit a great range of lifestyles (solitary and colonial), trophic diversity (heterotrophic or mixotrophic bearing photosynthetic symbionts) and tremendous morphological variability, spanning a wide range of sizes, from micrometers to several millimeters [2, 3]. Among the phylogenetic groups of Radiolaria, Acantharia [4] is the sole group of Radiolaria bearing a strontium sulfate skeleton, while the Polycystines group possesses opaline silica skeletons [5]. In the Polycystines group, Spumellaria [6] exhibits concentric and spherical shapes, while Nassellaria [7] showcases a diverse range of features. These include conical skeletons and a limited number of spicules, or even large colonies, akin to that found in the Collodaria group [8], or large individual skeletons as found in the Orodaria group [9]. This extensive diversity makes Radiolaria an ideal lineage for investigating adaptive evolution in protists.

Additionally, Radiolaria have successfully adapted to diverse ocean environments and are found at various depths, being important contributors to contemporary biogeochemical cycles. They can participate in the transformation of sinking particles and have been identified as a critical source of organic carbon [10, 11]. Global estimates indicate that Rhizaria, including Collodaria among other protists, may constitute up to 20% of the biomass of the euphotic layer (0-200m) [12]. They also play a substantial role in zooplankton abundance, contributing up to 33% [1]. In some areas, Radiolaria may dominate the eukaryotic community recovered from metabarcoding analyses of sediment traps based below the euphotic layer [13, 14]. They also have an essential role in the silicon cycle, as many representatives take up silicic acid from the seawater to form their skeleton [15] contributing approximately 20% to the biogenic silica production of the global ocean [16]. It is because of their intricate mineral skeleton made of opaline silica that Radiolaria have left behind a continuous fossil record since the Cambrian, more than 500 million years ago (Mya) [17], and have been instrumental for paleoenvironmental reconstructions [18, 19].

Due to the challenges of collecting these protists intact, the difficulty of maintaining them in culture, and the lack of information about their genomes, most of our current understanding about their ecology is derived from isolated and random observations in the field. Notably, various types of trophic modes and feeding strategies have been observed in Radiolaria. Some feed upon sinking particles while others are active predators or exhibit a combination of both behaviors. They use a network of cytoplasmic extensions (i.e., pseudopods) to capture and entangle a wide range of prey, from bacteria to multicellular heterotrophs like nauplii larvae [2, 20], even in oligotrophic waters where prey abundance is scarce. Many are mixotrophs harboring photosynthetic algae [21–23]. These symbionts can be found either directly inside the cell membrane as in Acantharia [24], or scattered on a gelatinous organic matrix surrounding the skeleton as in Collodaria [25]. The presence of photosymbionts in these organisms contributes to primary production. In the upper 20 m of the central North Pacific, Acantharia alone can contribute up to 4% of total primary production and up to 20% of surface production [26, 27]. It has been suggested that such symbiotic adaptations were triggered by long periods of accentuated oceanic oligotrophy in the Jurassic (ca. 175-93 Mya; [4]).

In contemporary oceans, mixotrophy allows Radiolaria to thrive in various environments. Collodaria are an important component of zooplankton biomass in the epipelagic and are most abundant in tropical latitudes and nutrient-impoverished waters [28]. Conversely, other Radiolaria, such as Acantharia or Spumellaria, are better adapted to meso- and eutrophic waters, richer in nutrients. These differences in biogeographical patterns have been attributed to the coloniality of Collodaria in which all individual cells share nutrients among the colony through the gelatinous matrix [25].

The presence of the gelatinous envelope surrounding the cell [29] could be a crucial advantage in enabling them to survive and thrive abundantly in oligotrophic waters, where prey are scarce. The gelatinous matrix also provides a conducive environment for hosting symbionts while facilitating resource optimization and adaptation strategies that contribute to the survival and ecological success of the cell. Indeed, other marine organisms, such as *Phaeocystis* algae, can also form an extracellular matrix that can host bacteria [30, 31], providing a nutrient-rich and protective environment and allowing them to form blooms and dominate the planktonic community.

In an oligotrophic environment, we collected several Nassellaria specimens embedded in a gelatinous matrix. To our knowledge, only one other study has documented these specimens, originally described as *Sethoconus myxobrachia* [32], yet recently reassigned to the genus *Phlebarachnium* [33]. The study of these rare specimens offers a unique opportunity to investigate the presence of the gelatinous matrix, a trait that may be vital for adaptation to new environmental conditions, allowing us to formulate new hypotheses to better understand evolutionary processes in protists. We present here findings on the phylogenetic relationships and global biogeography of *Phlebarachnium*. Our approach involved sequencing rDNA genes, characterisation of the surrounding waters where these specimens were collected, and the analyses of metabarcoding datasets (Eubank; [34]). Here we hypothesize that the gelatinous matrix serves as an ecological adaptation to oligotrophy, as we observed it evolved independently in different groups of Nassellaria. This gelatinous matrix would increase cell volume, facilitating the capture and storage of scarce prey and nutrients. Additionally, it offers larger cellular space to host an increased number of symbionts providing an optimal environment for them, here identified within the genus *Scrippsiella*.

## Material and Methods

### Sample collection

Samples were collected during the P2107 California Current Ecosystem Long Term Ecological Research (CCE-LTER) cruise in the summer of 2021 aboard the R/V *Roger Revelle* (Fig. 1A). The sampling strategy was designed around 3 multi-day quasi-lagrangian cycles. Each cycle involved following the drift trajectories of experimental arrays, aiming to sample the same water mass. Individual casts of Conductivity, Temperature and Depth (CTD) were conducted at each station (∼about 10 per cycle). Discrete bottle samples were collected at various depths from each station to measure nutrients (silicic acid (Si(OH)_4_) and nitrate (NO_3_^-^)). Water column nutrient samples were measured following the standard operating procedures from the CCE-LTER. Briefly, macronutrient samples were collected and filtered through a 0.2-μm capsule filter before being frozen at -20°C for on-shore autoanalyzer analysis.

**Fig. 1.**
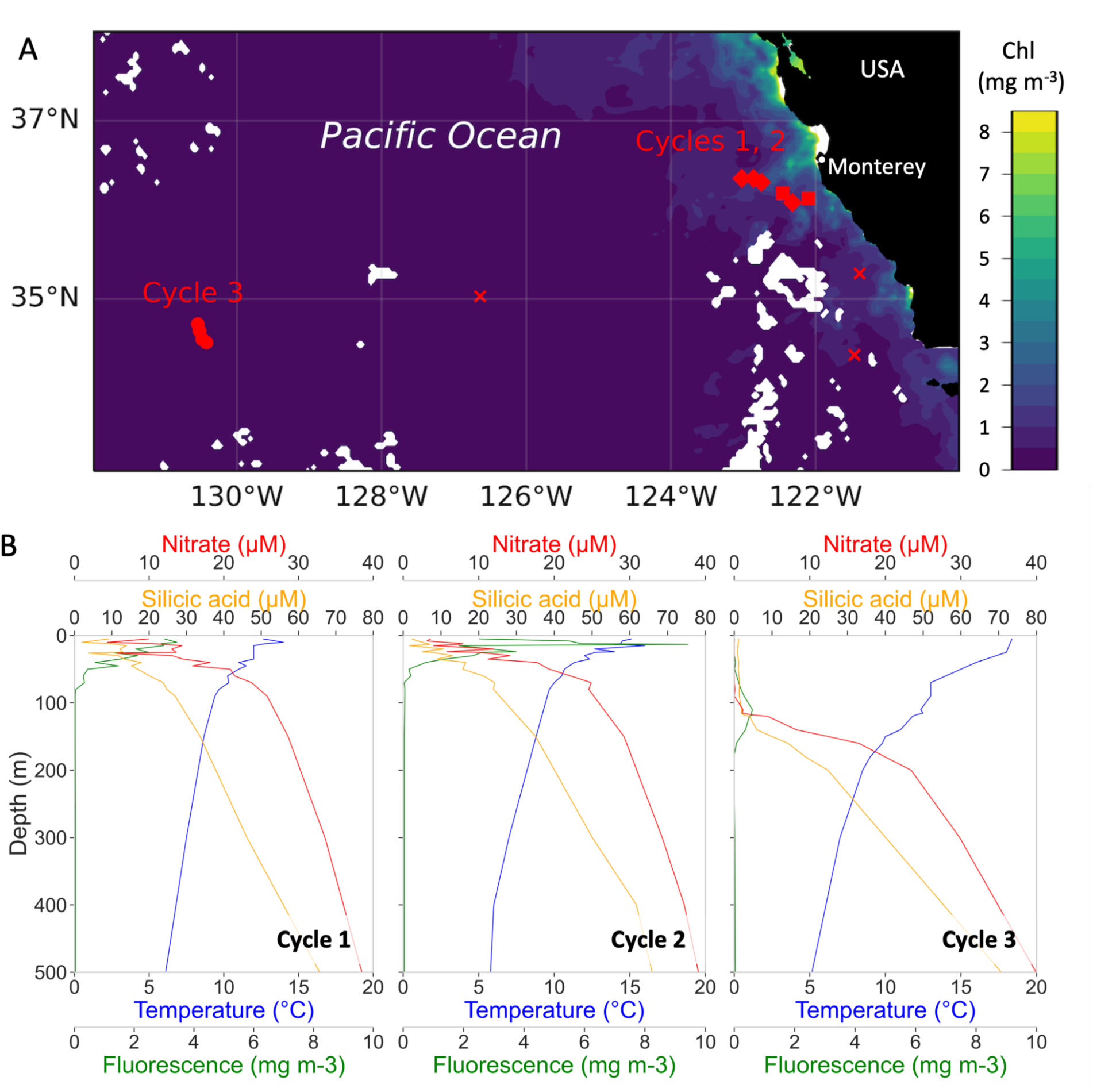
**(A)** Map of the sampling locations of the P2107 cruise where net tows were carried out. Squares are used for Cycle 1, diamonds for Cycle 2, circles for Cycle 3 and crosses for sampling outside the cycles. Cycle 3 stations are where Nassellaria specimens used in this study were found. The color scale shows real time Aqua MODIS chlorophyll concentration (mg m^-3^), 4 km resolution for August 2021. **(B)** Fluorescence (mg m^-3^) and temperature (°C) data from the CTD cast and nutrients concentration (silicic acid and nitrate, in µM) for each cycle.

In this study, we present results from a subset of 4 stations of Cycle 3 (3-6 August), sampled in oligotrophic waters, where Nassellaria identified as *Phlebarachnium* sp. according to [32] were found (**Sup Fig. S1**). Plankton samples were collected using a ringnet (200-µm mesh size, mouth diameter: 1 m) towed vertically from 50 m to the surface, from 300 m to the surface and from vertically stratified sampling collected with a 1-m^2^ opening, 202-µm mesh MOCNESS.

Immediately after collection, each net sample was observed using a stereomicroscope (Zeiss Stemi 508) and images of targeted organisms were captured using a Canon Eos 77D camera. Morphological measurements, including major and minor axes of individual cells, and their skeleton area, were obtained using ImageJ software [35, 36] to estimate the biovolume of the cells (Equations 1 and 2) (**Sup Table S1**). Each specimen was transferred into 1.5-ml Eppendorf tubes containing 50 μl of molecular grade absolute ethanol and stored at -20°C until DNA extraction. Scanning electron microscopy images were captured using a S-3400N Hitachi scanning electron microscope following the protocol in [37] for detailed examination of morphology.

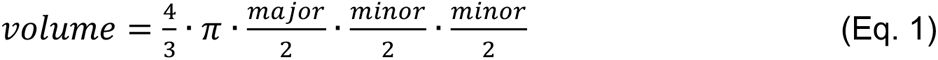

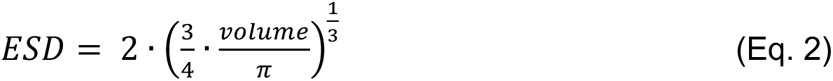

### DNA extraction and sequencing

DNA was extracted using the MasterPure Complete DNA and RNA Purification Kit (Epicentre) following the manufacturer’s instructions. Partial 28S rDNA (D1 and D2 regions) was amplified by Polymerase Chain Reaction (PCR) using Radiolaria and Nassellaria specific primers (28S-Nas-F: 5’ - AGT AAC GGC GAG TGA AGC - 3’ and 28S-Nas-R: 5’ - CCA ACA TAC DTG CTC TTG T - 3’). For further details about rDNA amplification see {Citation}. PCR amplicons were visualized on 1% agarose gel stained with ethidium bromide. Positive reactions were purified using the Nucleospin Gel and PCR Clean up kit (Macherey Nagel), following manufacturer’s instructions and sent to Macrogen Europe for sequencing. After sequencing, forward and reverse sequences were checked and assembled using ChromasPro software version 2.1.4 (2017). Sequences were compared to the GenBank reference database using the BLAST search tool integrated in ChromasPro to discriminate radiolarian sequences from possible contamination. The presence of chimeras was checked by mothur v1.39.3 [38] against the PR2 database v5.0. [39]. No chimeras were found, and therefore sequences were kept for further analyses. Final sequences were deposited in NCBI under the accession numbers OR978667-OR978672. Single-cell amplification and sequencing of the 18S rDNA was also attempted following [7], yet after checking correct *in silico* specificity of the primers, no successful reactions yielded 18S sequences of the Nassellaria host and DNA template was spent before achieving successful amplification.

### Phylogenetic analysis

Our partial 28S rDNA sequences were compared against an environmental dataset [40] containing near full-length sequences of the rDNA (from doi: 10.6084/m9.figshare.15164772.v3) using the ‘--usearch_global’ option from vsearch v2.21.1 [41]. In total 2 sequences of the near full-length rDNA were identified with a similarity identity of 99.8%, with the third best identity score at 89.1%. Additionally, 135 Nassellaria sequences were extracted from [42], with 24 Spumellaria sequences as the outgroup (**Sup Table S2**). Datasets of the 18S and 28S rDNA were independently aligned with mafft [43] using the L-INS-i algorithm (‘—localpair’) and 1000 iterative refinement cycles. The alignment was visualized in AliView v1.28 [44] to check for possible artifacts and trimmed with a 5% gap threshold using trimAl [45]. Final alignments contained 151 sequences with 1849 aligned positions for the 18S rDNA and 93 sequences with 800 positions for the partial 28S rDNA. Final datasets of the 18S and the partial 28S rDNA were concatenated with an in-house script freely available on GitHub (<u>MiguelMSandin/random/fasta/fastaConcat.py</u>), resulting in 165 sequences, 2647 aligned positions. Phylogenetic analyses were performed using Maximum Likelihood (ML) implemented in RAxML-ng [46] and IQ-Tree [47] and a Bayesian Inference (BI) implemented in MrBayes [48]. RAxML-ng was run under a GTR + Gamma model of evolution over 100 parsimonious starting trees and 1000 Felsenstein bootstraps. Prior to phylogenetic inference with IQ-Tree, ModelFinder identified GTR + F + R4 as the best fitting model of evolution and it was therefore used over 1000 bootstraps and 10 independent runs. Lastly, MrBayes was run under a GTR + Gamma model of evolution over 10 million generations and 4 independent runs. After checking for convergence of the bayesian MCMC chain in tracer v1.7.2 a 25% burnin was applied and a strict consensus tree was built.

While attempting 18S rDNA amplification, up to 6 different and high-quality sequences taxonomically assigned to dinoflagellates were sequenced from 6 independent hosts (deposited in NCBI under the accession numbers OR961112-OR961117). These sequences were combined with those used in [21] in order to study the phylogenetic relationships of the potential symbionts. In total 71 sequences of the 18S rDNA were aligned and trimmed (following methods stated above) resulting in 1803 positions. Phylogenetic analyses were inferred in RAxML-ng and IQ-Tree as mentioned above.

### Calibration of the molecular clock

Divergence times among sequences were estimated in BEAST v2.7.3 [49] under the Optimized Relaxed Clock [50] model to avoid assumption of substitution rates correlation between sister clades, a GTR+Gamma (4 categories) model of nucleotide substitution and a Birth-Death model of speciation [51]. Fossil calibrations were taken from [42], including a maximum root age of 1200 Mya and minimum of 500 Mya under a uniform distribution. The Bayesian MCMC chain was run over 10 million generations sampled every 1000 states on 3 independent runs (remaining operators were left as default). After completion, effective sampling size was checked to be equal or higher than 200 and the 3 independent runs were combined in LogCombiner v2.7.4 with a 25% burnin. The final tree was summarized in TreeAnnotator v2.7.3.

### Ancestral state reconstruction

We reconstructed ancestral states for the Nassellaria clade to infer whether the gelatinous matrix is an acquired trait or instead an ancestral trait. To do so, two complementary approaches were carried out over the resulting time calibrated tree. The first approach was performed under a Maximum Likelihood (ML) Binary State Speciation and Extinction model (BiSSE) implemented in the R package ‘diversitree’ v0.10 [52], with the functions ‘make.bisse’ and ‘find.mle’. Phylogenetic tips known for the presence or absence of gelatinous matrix [33] received a ‘1’ or a ‘0’, respectively, while environmental sequences with unknown morphology were assigned an empty value (i.e., ‘Not Available’: ‘NA’). All families described with the presence of a gelatinous matrix in [33] were represented in our dataset. The second approach followed a Bayesian Inference (BI) implemented in BEAST2 performed under a birth death model with 10 million generations sampled every 1000 steps and a 25% burnin. As for the ML approach, phylogenetic tips whose presence or absence of gelatinous matrix are known were given their respective categorical values. However, this approach does not allow the absence of character traits and thus those phylogenetic tips whose morphology is unknown were given a third categorical value. After such a limitation, the ML approach was chosen as principal analyses and the BI approach was used for assessing consistency of results and our dataset.

### Global biogeographic analyses

Global biogeographic patterns of *Phlebarachnium* were examined in the EukBank metabarcoding dataset of the V4 hypervariable region of the 18S rDNA [34] accessed at doi: 10.5281/zenodo.7804946. *Phlebarachnium* metabarcodes were extracted from the near-full length rDNA sequences mentioned above with the ‘--usearch_global’ option from vsearch v2.21.1 [41] and default parameters. In total, 2 metabarcodes from the V4 rDNA were selected and extracted from the EukBank dataset with a similarity identity of 99.5% and 96.9% (the third closest metabarcode hit to our *Phlebarachnium* sequences had an identity of 89.4%). Given that the abundance of the second metabarcode was significantly lower than that of the first metabarcode (14 total reads compared to 306522), read abundances of both metabarcodes were clustered together for biogeographic patterns exploration. Only samples with a marine identifier under ‘biome’ were considered in this study.

### Co-occurrence networks

To understand whether the presence of the gelatinous matrix has ecological implications or not, we reconstructed co-occurrence networks for all Radiolaria amplicons worldwide from the previously described dataset (EukBank). Network inference was performed using FlashWeave v0.19 [53], a recently developed tool to accommodate data compositionality and infer statistical co-occurrence or co-abundance interactions. All Radiolaria amplicons with known presence or absence of a gelatinous matrix were extracted from the OTU table, and samples with no Radiolaria reads were discarded. To avoid potential over-clusterization of intragenomic variability, PCR/sequencing errors and other artifacts [54], we only considered amplicons with a maximum relative abundance in any given sample of at least 1%. FlashWeave applies a centered log-ratio (CLR) transformation prior to inferring local-to-global learning framework, and thus no additional transformations were used. Given the relatively large proportion of amplicons assigned to undescribed groups of Radiolaria (such as RAD-A, RAD-B, RAD-C or environmental clades within e.g., Collodaria), analyses were then replicated also considering amplicons whose gelatinous state is unknown. In addition, given the large size ranges of gelatinous Radiolaria (from few µm to several cm), supplementary analyses were run on specific size fractions in order to account for possible sampling biases. In total, 945 Radiolaria amplicons were considered of which 244 were assigned to Radiolaria with a gelatinous matrix, 319 to Radiolaria without a gelatinous matrix and 382 had an unknown attribute. From the 563 amplicons with known attributes, 547 were found only in the small size fraction (0.22-20 µm), 501 in the medium size fraction (3-200 µm), and 495 in the large size fraction (80-2000 µm or bigger). Such overlap between different size-fractions groups was a result of the large variability of sampling approaches covered across the EukBank dataset.

Two additional analyses were applied over the resulting networks to better understand community structure and co-occurrence: The assortativity of the gelatinous attribute and Louvain clusters. The assortativity coefficient measures the preference of nodes with a given attribute to be connected to other nodes with the same attribute, and it was calculated using the function ‘attribute_assortativity_coefficient’ from the Python package ‘networkx’ v2.2 [55] implemented in the custom script <u>netAssortativity.py</u> (publicly accessible at <u>github.com/MiguelMSandin/random/networks/</u>). To test the significance of the assortativity values and the probabilities of deviating from random assortativities, we randomize the gelatinous attributes 100 times over the exact same network and compare the original assortativity value with those obtained from randomization with a one-sample T-test. The second analysis, the Louvain method, implements the multi-level modularity optimization algorithm for finding community structure in a hierarchical approach to extract non-overlapping communities or clusters. Louvain clusters were estimated with the ‘cluster_louvain’ function from the R package ‘igraph’ v2.0.3 [56] and a 0.5 resolution parameter since it was the value that best explained community structure by keeping small clusters yet sufficiently large to be meaningful regarding our initial question. All networks were visualized in Cytoscape v3.10.2 [57] after applying an organic layout.

## Results

### Environmental context

Specimens of *Phlebarachnium* were found in highly oligotrophic waters, where Chlorophyll *a* concentration was low (ranging from 0 to 9 mg m^-3^) (Fig. 1A). At Cycle 3, fluorescence values were almost negligible, except for a peak detected at around 100 m (Fig. 1B). In Cycles 1 and 2, near the coast, fluorescence and Chlorophyll *a* values were much higher in the upper 100 m of the water column than in Cycle 3. Nutrient concentrations (silicic acid and nitrate) in Cycles 1 and 2 showed an upward trend with depth from the surface waters. During Cycle 3, nutrients depletion was observed in the upper 150 m of the water column. Beyond this depth, values reached 30 and 25 µM for silicic acid and nitrate, respectively.

### Morphological assessment

Based on live observations and light microscope images, two populations of *Phlebarachnium* cells were defined showing differences in average size, color and depth distribution, which we refer to as the yellow and orange populations (Fig. 2A and B, respectively). Both populations were surrounded by a gelatinous matrix. Morphometric assessments of matrix and skeleton, including major and minor axes measurements, volume and estimated spherical diameter (ESD) calculations, were conducted on 25 individuals from each population (**Sup Table S1**). The ESD of the gelatinous matrix was 845 (± 73) µm and 1383 (± 129) µm, and the skeleton ESD was 332 (± 35) µm and 563 (± 49) µm for the yellow and orange populations, respectively. A Wilcoxon test indicated that both matrix and skeleton in orange specimens were significantly larger than yellow specimens (Z=-4.37, pvalue<0.001; Z=-4.37, pvalue<0.001). The skeleton volume/matrix ratio volume varied from 3 to 12%, with no significant differences among populations (Z=-1.70, pvalue=0.09).

**Fig. 2.**
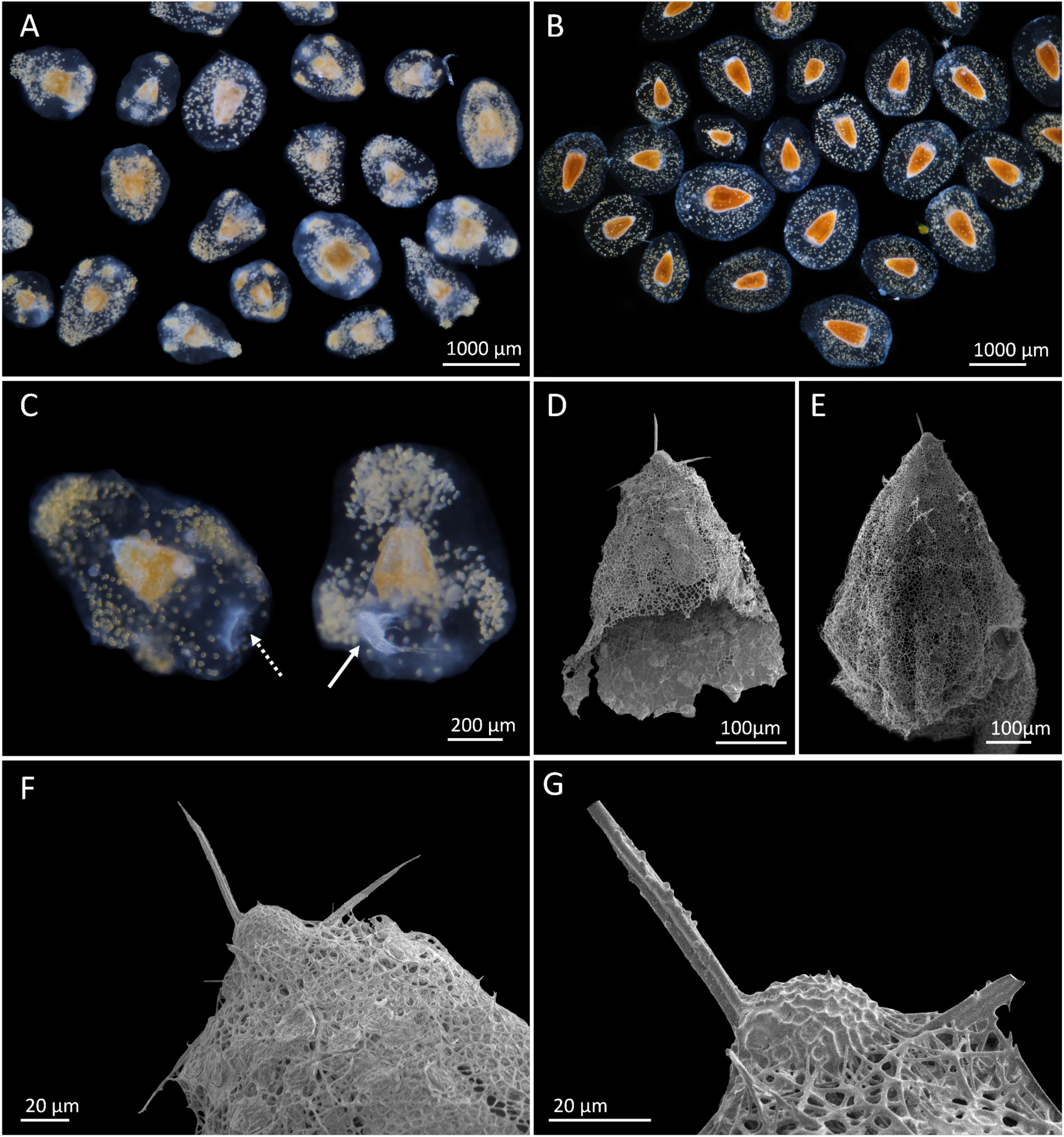
**(A-C)** Light microscope images of living *Phlebarachnium* specimens; **C** is a composition of two separate images, with the dotted arrow indicating the aperture within the gelatinous matrix, and the solid arrow pointing to the prey embedded in the matrix, **(D-E)** scanning electron micrograph of the skeleton of *Phlebarachnium* specimens, with a detailed focus of the cephalis **(F-G)** used for traditional morphological classification. **A, C** and **E** are the “yellow” population of *Phlebarachnium*. collected with the 0-50 m net tow; **B, C** and F are the “orange” population found below 75 m.

In approximately 15% of the specimens from the yellow population collected, we observed the presence of enclosed prey within the gelatinous matrix. These include mainly harpacticoid copepods and nauplius larvae. All copepods were consistently placed near the aperture (Fig. 2C, dashed white arrow), and on many occasions we observed a clear opening within the gelatinous matrix (Fig. 2C, solid white arrow). Although we cannot conclude on the vertical distribution of these organisms due to sampling limitations, we observed some distribution patterns in depth. While host specimens of the yellow population were found in all the nets tows, from 0-50 m, from 75-150 m and 0-300 m, specimens of the orange population were only found in the deeper net tows, below 75 m.

Both populations show numerous golden-yellow bodies distributed across the gelatinous matrix, with a lower concentration in the deeper-dwelling orange population. Both populations show different distributions of these golden yellow bodies. In the yellow population, they appear arranged in clusters within specific and recurring regions of the matrix, which may show rounded inflations. We noted the presence of 3 to 4 clusters of symbionts, with one positioned toward the apical horn and 2 to 3 arranged toward the bottom of the gelatinous matrix (**Fig. 2A**). In contrast, in the orange population, symbionts are evenly distributed throughout the matrix (**Fig. 2B**). Despite differences in skeleton size and symbionts distribution, both populations shared common features in their skeletal morphology. Each representative exhibited a reticulate shell, characterized by a spherical cephalic region and a larger bell-shaped thorax (**Fig. 2D-G**).

### Molecular and phylogenetic analyses

Phylogenetic analyses revealed that *Phlebarachnium* constitute a highly supported novel clade (Bootstrap Support, BS≥94; Posterior Probability, PP≥0.91) within Nassellaria and confidently supported (BS=72 and 78, PP=0.98) as sister to Lithochytridoidea, Pterocorythoidea and an environmental clade (a clade with no morphologically described sequences) named as Nass-4 (**Fig. 3**).

**Fig. 3.**
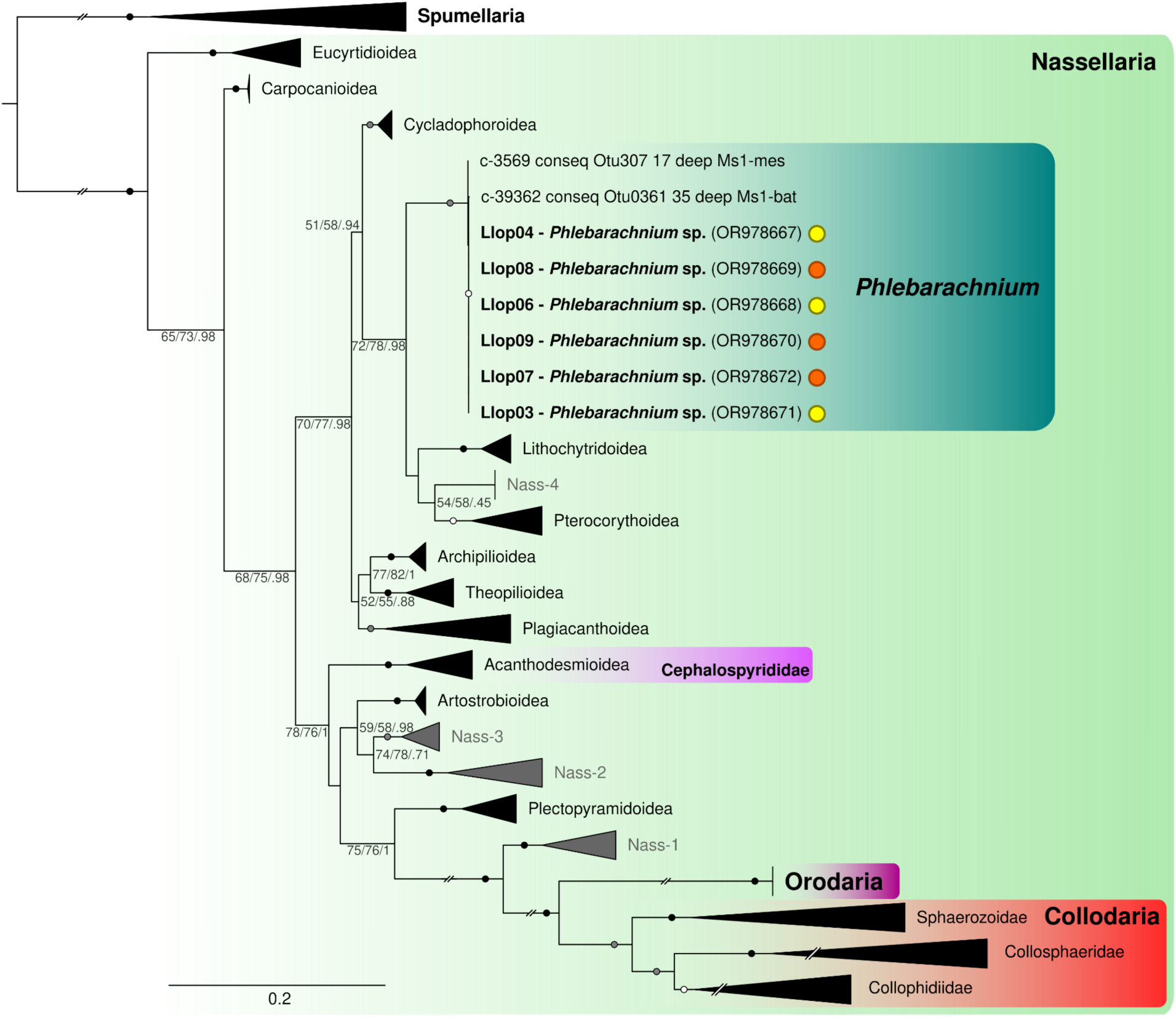
Molecular phylogeny of Nassellaria inferred from the concatenated 18S and the partial 28S (D1+D2) rDNA (141 taxa and 2649 aligned positions), highlighting all described groups with a gelatinous matrix. Note that within Acanthodesmioidea, only one family has been reported to be surrounded by a gelatinous matrix (represented in our dataset by *Ceratospyris* and *Lophospyris).* In total, 24 Spumellaria sequences were assembled as an outgroup. Values at nodes represent RAxML-ng (GTR+Gamma; tree shown in figure), IQ-Tree (GTR+F+F4; 10 runs) bootstraps values (BS; 1000 bootstrap each) and MrBayes (GTR+G) posterior probabilities (PP; computed after 10 million generations over 4 independent runs and 25% burnin). Only values above 50 BS and/or 0.5 PP are shown. Hollow circles at nodes indicate BS ≥ 80 and PP ≥ 0.8, filled grey circles BS ≥ 90 and PP ≥ 0.9, and filled black circles BS ≥ 99 and PP ≥ 0.99. Branches and clades with a double barred symbol have been 50% reduced for clarity. Orange and yellow circles after the *Phlebarachnium* sequence names represent whether they belong to the so-called “orange” or “yellow” populations.

This distinct *Phlebarachnium* clade shows a distant phylogenetic relatedness to Orodaria, Collodaria and the other family within Acanthodesmioidea that also develops a gelatinous matrix around the cells. Molecular dating estimates the last common ancestor between *Phlebarachnium* and Collodaria in 337 Mya (with a Highest Posterior Density -HPD-between 415 and 270 Mya; **Fig. 4**). Orodaria and Collodaria diversified 166 Mya (HPD: 223-115). While Collodaria has been dated to first diversified at 134 Mya (HPD: 185-92), *Phlebarachnium* clade first diversified 5.4 Mya (HPD: 13.8-1.1), yet separated from its sister clades much earlier at 127 Mya (HPD: 187-84), and Orodaria only diversified 1.3 Mya (HPD: 6.4-0.01).

**Figure 4.**
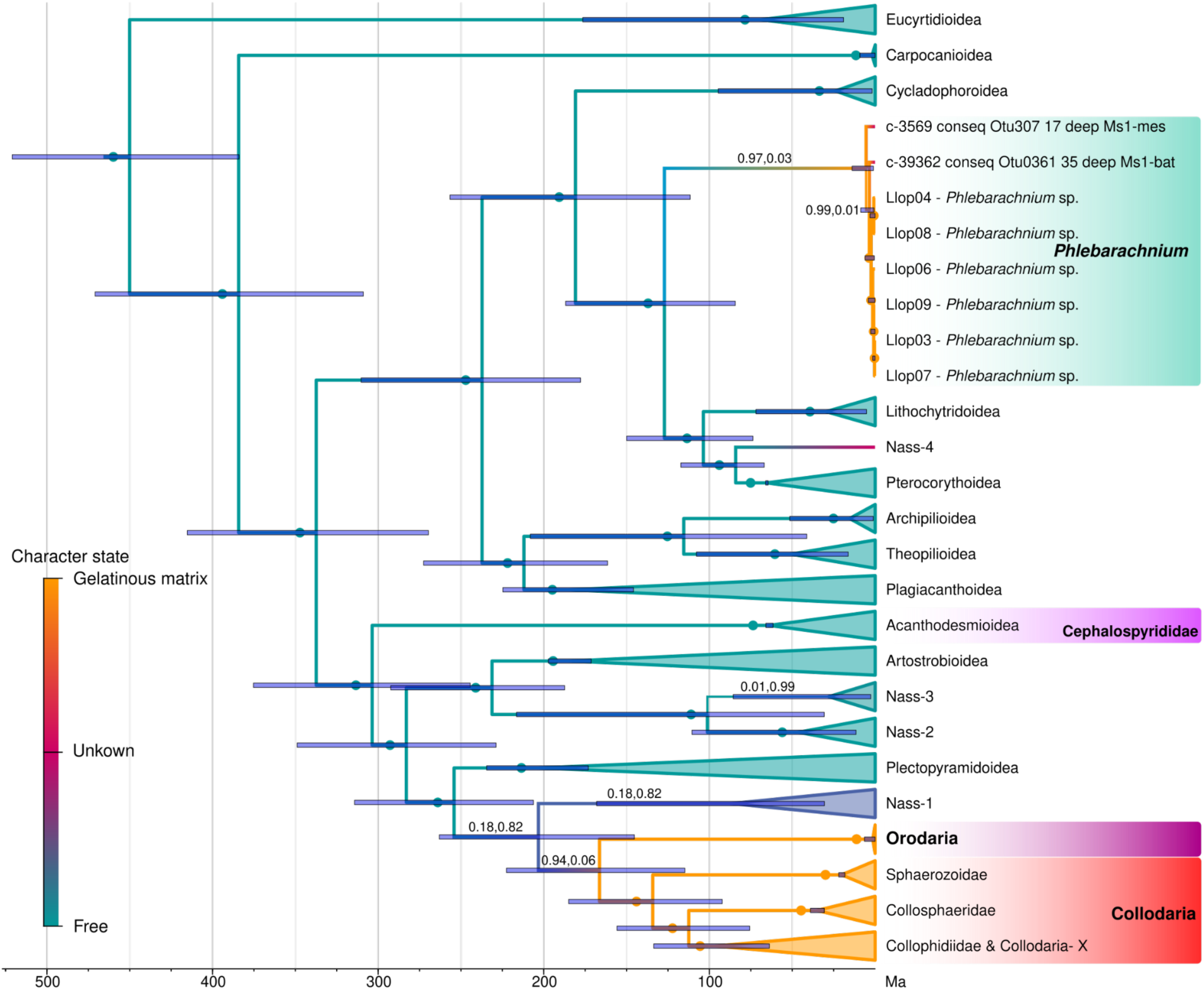
Time-calibrated tree (Molecular clock) of Nassellaria highlighting in the branch color the ancestral state reconstruction probabilities between a state with presence of a gelatinous matrix or absence (free). Divergence times were inferred under an optimized relaxed clock model and GTR+G for 3 independent chains of 100M cycles sampled every 1000 steps and a 25% burnin, implemented in the software package BEAST2, on the alignment matrix used for phylogenetic analyses (Fig. 3). Twelve different nodes were selected for the calibration, based on [40] with the exception of the root in which a uniform distribution (with minimum of 500 and maximum of 1200 Mya) was used. Ancestral state reconstruction was performed under a binary state speciation and extinction model (BiSSE) implemented in the R package *diversitree*. Values above branches represent the probability of a gelatinous ancestral state and of the absence of a gelatinous matrix respectively. When full support for either state, probabilities are represented by a dot. Blue bars indicate the 95% highest posterior density (HPD) intervals of the posterior probability distribution of node ages.

Ancestral state reconstruction analyses confidently support an independent development of the gelatinous matrix in *Phlebarachnium*, the group of Collodaria and Orodaria, and the family Cephalospyrididae within Acanthodesmioidea (**Fig. 4**). The reconstructed last common ancestor between all gelatinous groups had no gelatinous matrix and such state was fully supported in both maximum likelihood and Bayesian inference. However, given the amount of undescribed diversity (especially within Collodaria), Bayesian inference failed to reconstruct a state for the last common ancestor between Orodaria and Collodaria (**Sup Fig. S2**) with 0.48 Posterior Probabilities (PP) of an unknown state over 0.47 PP for a gelatinous state and 0.05 PP for a non-gelatinous state. The gelatinous matrix could have evolved independently in Collodaria and Orodaria, and thus the first appearance of the gelatinous matrix could have either be with the diversification of Orodaria and Collodaria (at ∼166 Mya; or earlier) or with that of Collodaria (at ∼134 Mya), nearly matching the divergence of *Phlebarachnium* with their sister groups (at ∼127 Mya).

While morphological observations identified two distinct populations, molecular analyses showed no clear differentiation as all sequences obtained in our study were identical. Environmental molecular diversity from the different datasets studied herein also showed limited diversity, agreeing on a unique molecular marker of *Phlebarachnium*, despite the different lengths (near full-length rDNA) and sequencing depth (accounting for intragenomic variability) considered in the respective environmental datasets. In addition, both populations show similar golden yellow bodies phylogenetically assigned to *Scrippsiella* (**Sup Fig. S3**), a dinoflagellate frequently found in symbioses with other organisms, yet independent of Polycystines’s common symbiont, *Brandtodinium*.

### Biogeographic analyses

Overall, the collection and identification of this enigmatic Radiolaria in the California Current allowed for the identification of a previously unassigned sequence in a global metabarcoding dataset. Here, we have used different sequences coming from various sources to explore global biogeographic patterns of *Phlebarachnium*, and due to their high degree of similarity (>96.9%) and the pairwise distance dissimilarity to other non-*Phlebarachnium* sequences (<89.4) we were confidentially able to taxonomically assign these reads to *Phlebarachnium* (**Fig. 5D**). Global metabarcoding analysis of the V4 hypervariable region of the rDNA shows that *Phlebarachnium* contributed on average 0.45 (±1.4) % to the total eukaryotic community across the EukBank dataset (with a median contribution of 0.02%). Yet in some samples of the Pacific Ocean, *Phlebarachnium* contributes to up to 13.41% of the total eukaryotic reads (**Fig. 5A**). The Pacific Ocean accounted for the majority of *Phlebarachnium reads*, gathering more than 96% of the total *Phlebarachnium* relative reads and nearly half of the samples where *Phlebarachnium* was present (75 of 162 global samples of which in 58 showed a relative abundance of at least 2% to the total eukaryotic reads). The highest relative abundance of *Phlebarachnium* coincided with the oligotrophic gyres of the Pacific Ocean at ∼30° of latitude (**Fig. 5B**) and in between -80° and - 180° of longitude of the upper mesopelagic water mass (200-1000 m; **Fig. 5C**).

**Fig. 5.**
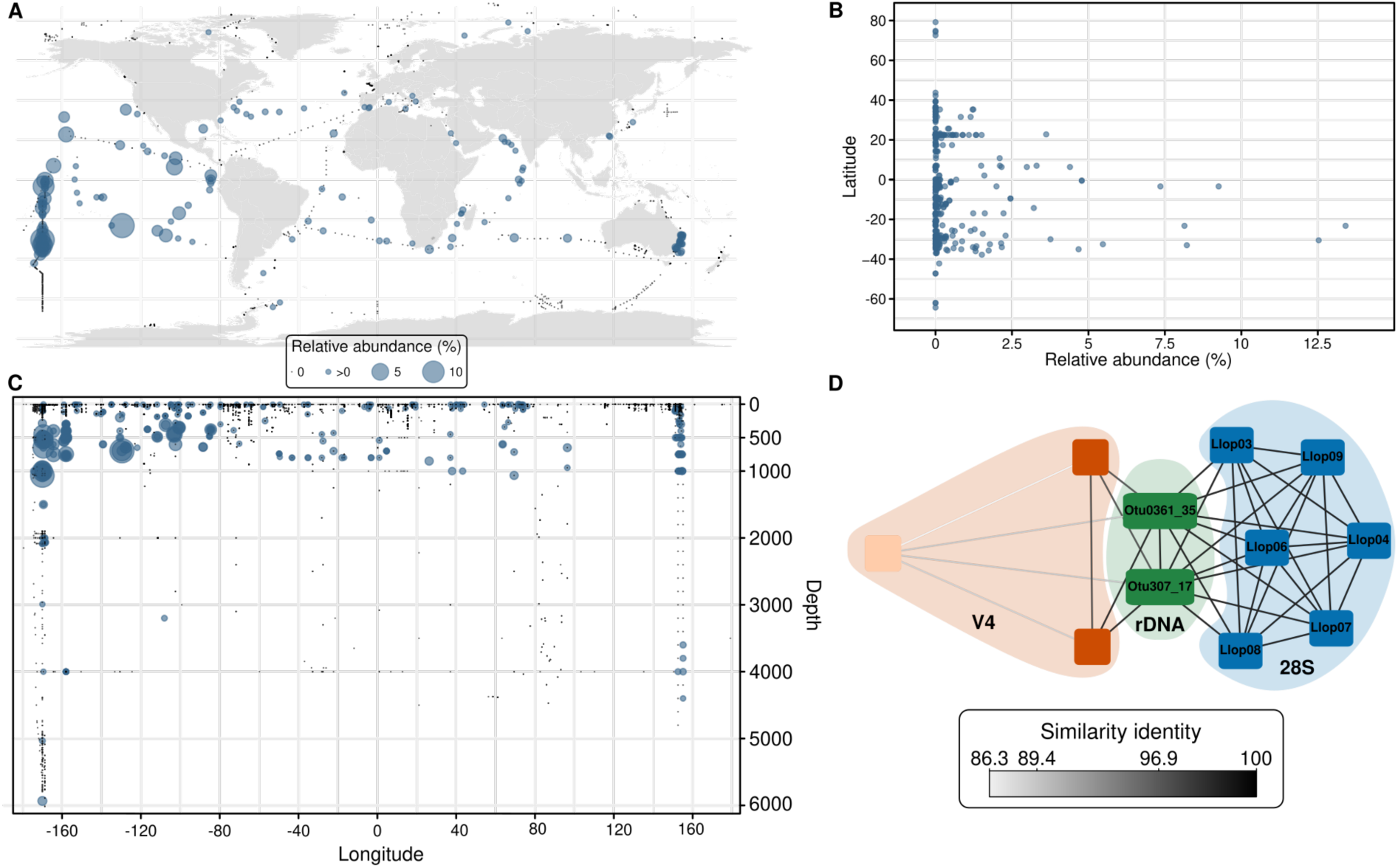
Global biogeography of *Phlebarachnium*. inferred from the EukBank dataset (Berney et al. 2023) on samples with a marine identifier. **(A)** World map showing the different sampling stations (black dots) and maximum contribution of *Phlebarachnium*. to the total eukaryotic community (blue dots) per locality. **(B)** Relative abundance of *Phlebarachnium*. across the latitudinal gradient. **(C)** Depth profile of *Phlebarachnium*. across the longitudinal gradient. Note panels A, B and C are aligned so that latitude and longitude can be cross-linked (a very dimmed grid can be seen in panel A for the main parallels and meridians extended from panels B and C respectively). **(D)** Graphical representation of the similarity identity (computed with vsearch: ‘allpairs_global’; and represented by the grey gradient lines, or edges) between all *Phlebarachnium* sequences (coloured squares, or nodes) used in this study. Blue nodes represent the D1+D2 regions of the 28S rDNA sequenced in this study, green nodes represent the near full-length rDNA, and orange represent the V4 hypervariable region of the 18S rDNA from eukBank [34]. Note the light orange node representing the third closest hit to our reference sequences, and confidentially showing the taxonomic assignment of *Phlebarachnium* used in this study. Node codes refer to the sequence identifier for referencing across figures (e.g. Fig. 3)).

Co-occurrence network analyses showed a strong correlation in the biogeography of gelatinous Radiolaria with other gelatinous Radiolaria and similarly for non-gelatinous Radiolaria co-occurring with non-gelatinous Radiolaria (**Fig. 6**). Such a pattern received an assortativity coefficient of 0.79, with a p-value of 0 (**Fig. 6A**). Louvain clusters identified three communities, of which two tended to contain amplicons that show a higher abundance towards the surface whereas the third community showed amplicons co-occurring at deep samples (**Fig. 6B**). Among the two shallow water communities one of them was constituted of primarily gelatinous Radiolaria and was correlated with no environmental parameter. The second shallow community contained primarily non-gelatinous Radiolaria and was positively correlated with nearly all environmental parameters, especially with total eukaryotic ASV richness, meaning that these amplicons tend to co-occur when samples are taxonomically diverse. The family Cephalospyrididae (Acanthodesmioidea) are the only amplicons associated with a gelatinous matrix that appeared in this cluster, and their maximum relative abundance is lower (with maximum relative abundances of 9.1, 4.4, 2.1, 1.6 and 1.2%) compared to other amplicons with a gelatinous matrix (e.g., *Phlebarachnium*: 13.41%). *Phlebarachnium* was in the third Louvain cluster and was correlated with depth. Despite that only 18 amplicons correspond to gelatinous Radiolaria (from 142 amplicons) in the third cluster, this attribute still showed a significant assortativity coefficient (0.27).

**Figure 6.**
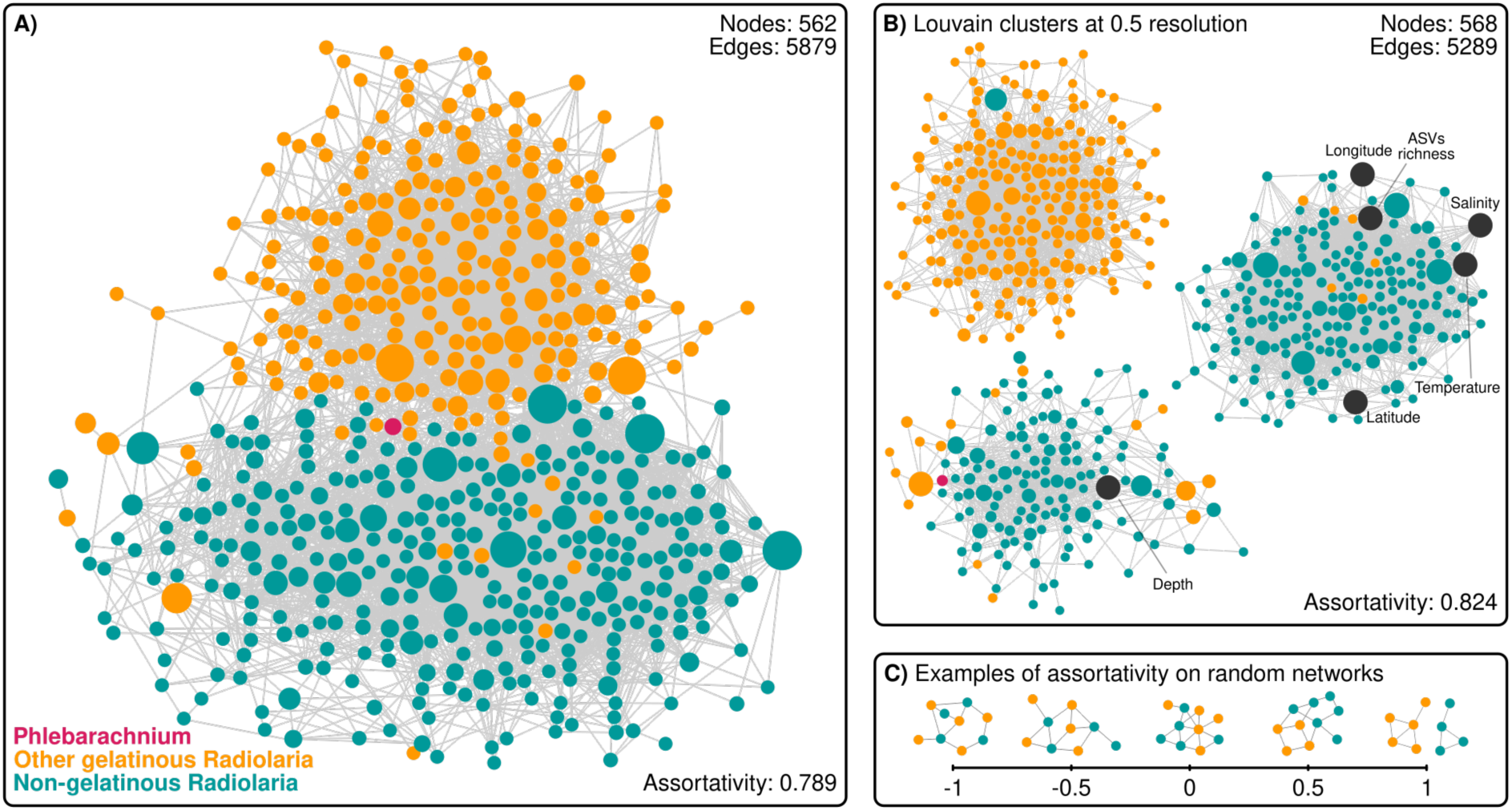
**A)** Co-occurrence network of Radiolaria across the global oceans highlighting gelatinous and non-gelatinous amplicons (nodes) extracted from the EukBank dataset. Co-occurrence was estimated in FlashWeave, including only amplicons with a maximum relative abundance (to any given sample) of at least 1%, in order to remove co-occurrence of sequencing errors and other artifacts, and amplicons whose gelatinous attribute is known (please see **SupMat Fig. S3**, for a detailed co-occurrence of different size fractions and **SupMat fig S4** for co-occurrence analysis of amplicons with both known and unknown attributes). For clarity, negative connections and environmental variables were removed from the graph. **B)** Louvain clusters of the original network shown in A) (including negative connections and environmental variables) explaining community structure and co-occurrence. Environmental variables are shown in dark gray circles. Size of the nodes represent the maximum relative abundance for visual guidance. **C)** Random networks with given attributes to exemplify different levels of assortativity.

When breaking down co-occurrence patterns into different size fractions to account for possible sampling biases, similar assortativity coefficients were found with 0.749, 0.646 and 0.734 coefficients for the small, medium and big size fractions respectively (**Sup Fig. S4**). In all different size fractions, Louvain clusters that contained *Phlebarachnium* were never found to be correlated with total ASV richness, and neither were other clusters containing primarily gelatinous amplicons. Besides, *Phlebarachnium* showed correlations with deep samples only in the small size fractions, while in the medium and big size fractions there is no such correlation. The inclusion of amplicons with an unknown trait also confirmed the trend of gelatinous amplicons co-occurring with gelatinous amplicons and non-gelatinous with non-gelatinous (**Sup Fig. S5**). Since an unknown attribute cannot be given, it represents a third attribute and yet a significant artificial assortativity coefficient of 0.298 (or 0.796 after removing the unknown trait) and three independent Louvain clusters were also identified as in **Fig. 6**.

Given the strong statistical support for the contrasting co-occurrence of amplicons with a gelatinous matrix and amplicons without a gelatinous matrix, we repeated these co-occurrence networks analyses including only samples from the *Tara* Oceans expedition and similar patterns were found (data not shown).

## Discussion

Here, we identified the genus *Phlebarachnium*, a distinct Nassellaria clade showing a characteristic gelatinous matrix surrounding the cells and highly abundant in oligotrophic waters. *Phlebarachnium* is, to our knowledge, the only solitary lineage of Radiolaria with a gelatinous matrix that has been reported to exhibit bloom-like growth in oligotrophic waters. Despite their distant relationship within the Nassellaria group, *Phlebarachnium*, Orodaria and Collodaria share the adaptation of producing a gelatinous matrix and affinity towards oligotrophic zones. Orodaria is restricted to oligotrophic deep waters, predominantly found in the Pacific Ocean, and lacks symbiotic algae within its gelatinous matrix [9]. Collodaria exhibits high abundance and diversity in shallow, oligotrophic waters [58], a trait attributed to the presence of numerous symbionts scattered in the gelatinous matrix of the colonies [28]. Other Radiolaria groups, such as Acantharia, do not possess a gelatinous matrix but still host numerous photosymbionts [24]. Since Acantharia are rarely found in oligotrophic waters, here we argue that the gelatinous matrix could be a specific innovation to cope with nutrient-poor systems. The gelatinous matrix would increase the volume-to-body weight ratio, facilitating the hosting of a larger number of symbionts and establishing a particular microenvironment that favours the viability of these organisms under conditions of nutrient depletion. Additionally, the increased volume-to-body weight ratio would improve the chances of prey encountering and/or scavenging nutrients for mineralization of the skeleton and/or symbiont nourishment. For non-photosymbiotic Orodaria, mostly found in the mesopelagic, the gelatinous matrix may play an important role for buoyancy. Altogether, the gelatinous matrix seems to be a convergent adaptation to oligotrophic waters, where other Radiolaria relatives are rare.

Other protists have also developed sophisticated behaviors to enhance encounter rates. Some dinoflagellates can build a mucosphere with the capacity to attract and immobilize diverse microbes, facilitating subsequent prey selection [59]. In other Radiolaria groups, comparable independent innovations in life mode and morphology have already been hypothesized and documented. Nassellaria’s closest relative, Spumellaria, experienced a substantial increase in planar forms during the second half of the Mesozoic, (ca. 145 Mya), which is thought to have resulted from low nutrient availability favoring an increase in the surface/volume ratio [8]. Here we have shown that the development of the gelatinous matrix happened at nearly the same geological time and we have attributed similar oligotrophic reasons. During the same geological epoch, Acantharia, established a photosymbiotic relationship with the haptophyte *Phaeocystis* also linked to extended periods of oligotrophy [4]. In such a relationship, photosymbionts of Acantharia exhibit plastidal volumes 38-fold larger than those of free-living *Phaeocystis* [24].

In the case of Acantharia, their photosymbionts are located inside the cytoplasm [24], whereas Collodaria harbors their photosymbionts scattered in the gelatinous matrix or located on the capsules. To our knowledge, Collodaria has not been documented without symbionts [58]. Moreover, some Collodaria have been shown to control the physiology and perhaps the life cycle of their photosymbionts to an extent [25]. In the yellow population of *Phlebarachnium* studied here, the symbionts were arranged forming discrete clusters within the host’s gelatinous matrix, suggesting an active host-mediated placement. In contrast, in the orange population found living at deeper water masses, the symbionts were evenly distributed throughout the matrix. Despite morphological variations among *Phlebarachnium* specimens, DNA barcoding and environmental molecular diversity showed almost identical sequences in all specimens studied here, with differences across the rDNA significantly smaller than those found at the intragenomic level [54]. This could also suggest that there exist various morphotypes in the same species. The co-occurrence network analyses showed a depth profile of *Phlebarachnium* where the smallest size-fractions correlate with deep samples, and larger size fractions tend to co-occur at the surface. It is believed that small bi-flagellated cells (called swarmers), released by mature cells, play a role in the reproductive cycle of Radiolaria [60, 61], and therefore such morphological plasticity and symbiont relocation might represent different life-stages. Regardless of morphological differences, all specimens of *Phlebarachnium* hosted round, golden-yellow bodies scattered in the gelatinous matrix. Molecular analyses identified such bodies as *Scrippsiella* (Peridiniales, Dinophycea), a common planktonic symbiont of the jellyfish *Velella velella* [62] and also seen in symbiosis with Acantharia [4], but never before reported in Polcystines Radiolaria [21, 22]. However, given the broad symbiotic interactions among the Dinophycea group and the long divergence time of *Phlebarachnium* (∼127 Mya to the last common ancestor of the closest relative), it is plausible that they developed a symbiotic relationship with a different group.

In addition to playing a central role in resource acquisition and harbouring symbionts, the matrix also could be important for prey capture. Nassellaria are known to have cytoplasmic extensions (pseudopods) that emerge from the conical cephalis, forming a web of cytoplasmic filaments where prey are captured [63]. The presence of the extracellular matrix likely provides the cell with greater motility and flexibility compared to cells lacking this matrix, facilitating predation on other cells. In many of the collected specimens, we observed the formation of a hole in the matrix, containing enclosed prey, predominantly copepods of the order Harpacticoida and nauplius larvae. Pelagic harpacticoids, such as *Microsetella* sp., are often associated with marine snow and aggregates in offshore areas where suspended food is scarce, similar to the waters where the *Phlebarachnium* specimens of this study were collected. The consistent occurrence of prey in the gelatinous matrix beneath the cephalis suggests that *Phlebarachnium* may play a relevant role in pelagic food webs by preying on metazoan taxa. This is particularly notable in oligotrophic waters, where *Phlebarachnium* seems to be endemic and prey are scarce.

In our study we found *Phlebarachnium* in high relative abundances in oligotrophic waters, even in specimens collected at deeper depths (0-300 m). Metabarcoding analyses show that *Phlebarachnium* reaches high relative abundances in the mesopelagic zone (200-1000 m), where light penetration is limited and possibly insufficient for the functionality of the symbionts’ photosynthesis. Taking into account the significant high relative abundance below 500 m around -160° longitude, this implies that these organisms may be versatile, thriving solely through heterotrophy without relying on photosynthesis. However, oligotrophic clear waters display relatively deep chlorophyll maxima (Fig. 1B) and thus photosynthesis is found at deeper environments regarding nutrient enriched waters. Collodaria has already been shown to inhabit ocean areas where other Radiolaria relatives are rare [42]. In tropical and subtropical nutrient-depleted surface waters, Collodaria can reach up to 1.6 x 10^4^ - 2.0 x 10^4^ colonies per m^3^ [64, 65]. In our study, *Phlebarachnium* sequences reached maximum abundances of 13.4% relative to the total eukaryotic community reads, especially towards the upper mesopelagic. In this context, *Phlebarachnium* likely thrives in deeper water layers than Collodaria, establishing a distinct ecological niche. The access to this environment allowed them to become important contributors to the planktonic food web of an ecosystem with potentially low predators. However, caution is required when interpreting metabarcoding data [66, 67]. Life cycle of Radiolaria remains poorly understood, and the same metabarcoding reads might reflect different life-stages (as previously discussed) or it might happen that dead cells are incorporated into sinking particles [68].

The high abundance and local endemism of *Phlebarachnium* to oligotrophic waters makes them useful candidates for paleoenvironmental reconstruction studies. Their bloom-like presence might reflect an important role in biogeochemical cycles. Considering climate change and expansion of oligotrophic waters in certain oceanic regions [69], the gelatinous matrix could be an enormous advantage for these organisms. Specializing in resource-poor waters could allow *Phlebarachnium* to expand into novel ecological niches, such as different habitats or trophic modes and adapt to novel environmental conditions. It has already been observed that ocean warming can trigger major shifts in the geographic distribution due to the combined effects of warming, stratification, light, nutrients and predation, in a wide range of organisms, from phytoplankton to mammals [70, 71]. In fact, the extensive fossil record of Radiolaria has already shown that rapid climate changes can lead to extinction of specific groups [72]. It is therefore critical to understand whether adaptations, such as the development of a gelatinous matrix, can help certain organisms cope with the rapid changes in ocean chemistry and temperature expected due to anthropogenic activity.

Our study highlights that the presence of a gelatinous matrix in *Phlebarachnium* could be an original strategy to cope with oligotrophy in the oceans. Studying the exact role of the matrix in these organisms is of particular importance to better understand biodiversity and adaptation of Radiolaria. In particular, investigating the genes responsible for its formation and its chemical composition (i.e., the amount of lipids and carbohydrates) may help to further understand its relationship with symbionts, prey and the host cell. However, both the genomes of Radiolaria and their reproduction in culture have so far withstood our efforts, which complicates further investigations with traditional or conventional techniques. That is why these observations, despite being random, are crucial, allowing us to hypothesize that the gelatinous matrix might be an advantageous trait to adapt to oligotrophic conditions by increasing the effective body size and creating an advantageous microenvironment for symbionts. Such increase in size might thus allow them to host more photo-symbionts than their relatives without a gelatinous matrix and similar body weight and further digest prey. This study not only advances our understanding in the ecological strategies of Radiolaria but also offers insights into the broader dynamics of organisms flourishing in nutrient-limited ecosystems.

## Author contributions

NLM and ML collected the samples. NLM and MMS conceived the presented idea, participated in the conceptualization, planned and performed the analyses, prepared the figures and wrote the manuscript. SR carried out the DNA extraction, amplification and purification. NLL, ML, SR, YN, TB and MMS commented and revised the manuscript.

## Acknowledgments

The authors would like to thank the captain and crew of the R/V *Roger Revelle* of the Scripps Institution of Oceanography for their expertise and support. We also thank colleagues within the CCE-LTER program (National Science Foundation grants OCE-1637632 and OCE-1614359) who contributed to sample collection and its lead PIs Katherine Barbeau and Mark Ohman. This work was supported by the “Agence Nationale de la Recherche” project RhiCycle (ANR-19-CE01-0006) granted to TB and the ISblue project RHICE (ANR-17-EURE-0015). NLL received funding from the European Union’s Horizon Europe research and innovation programme under grant agreement No 101064167 (MSCA postdoctoral fellowship Si-ORHIGENS). MMS was supported by a postdoctoral fellowship from the Beatriu de Pinós programme of the Government of Catalonia’s Secretariat for Universities and Research of the Generalitat de Catalunya Economy and Knowledge (grant number: 2021BP00068). We would like to express our gratitude to Aude Leynaert for support, Fabrice Not for providing lab space and resources, ABiMs for computational resources, Johan Renaudie for the taxonomic contribution, Nicolas Henry for advising us on biogeographic analyses, Ian Probert for discussions on the nature of the symbionts and Steve Cunningham for SEM support. We are also very grateful to Colleen Durkin and Daniel Richter for their valuable input and advice on earlier versions of this manuscript. We thank two anonymous reviewers for their valuable contributions to improving the manuscript.

## Data availability

The datasets generated during and/or analysed during the current study were deposited in NCBI and are available under the accession numbers OR978667-OR978672 and OR961112-OR961117. The environmental datasets used were accessed from doi: 10.6084/m9.figshare.15164772.v3 and <u>10.5281/zenodo.7804946.</u>

## Supplementary Material table headers

**Table S1**. Morphological measurements, major and minor axes of individual cells, and their skeleton and gelatinous matrix areas.

**Table S2**. List of specimens used for phylogenetic analysis.

**Figure S1.**
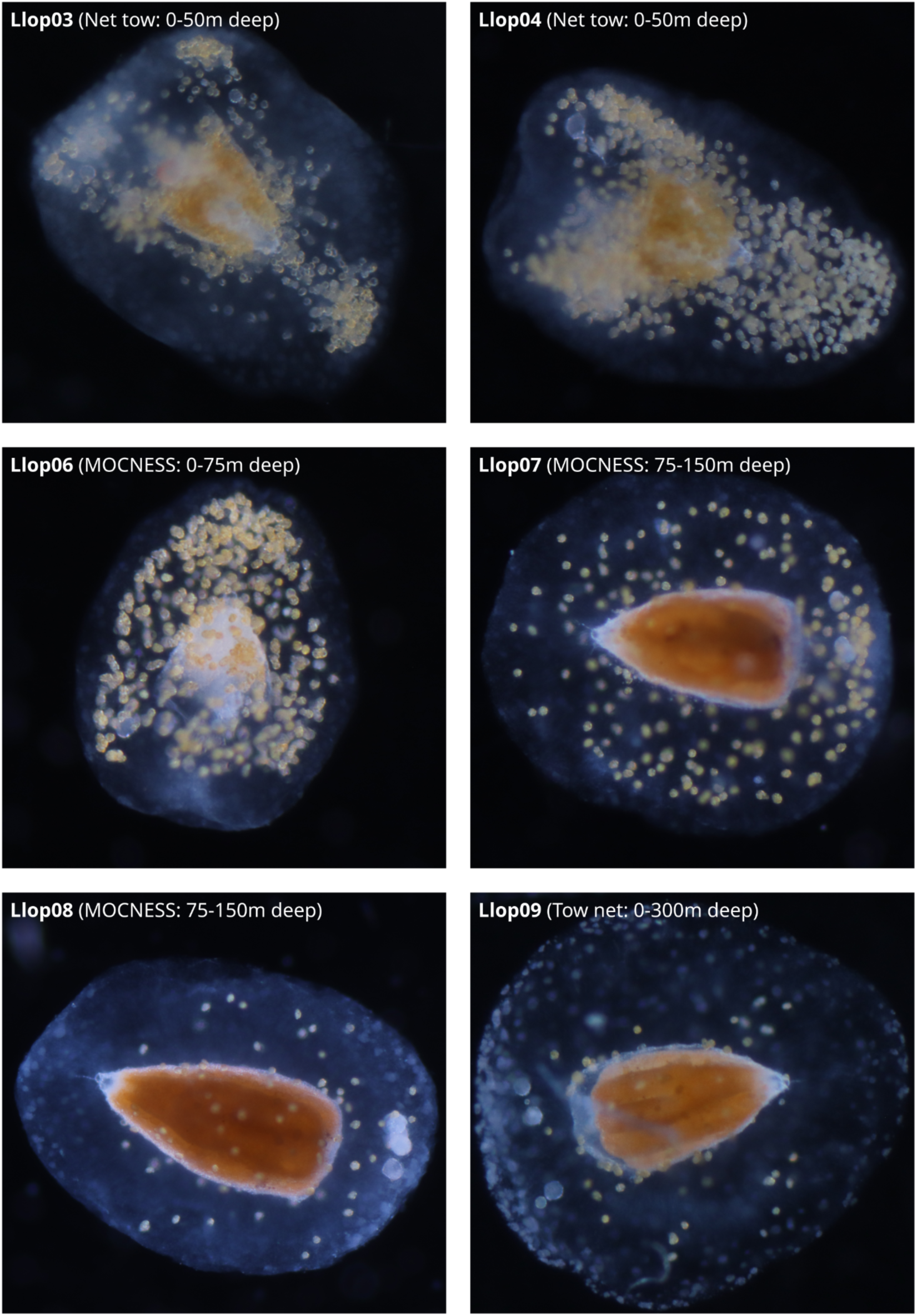
Light microscope images of live *Phlebarachnium* specimens used in this study for phylogenetic analysis in Fig. 3. Scale bar represents 500 μm.

**Figure S2.**
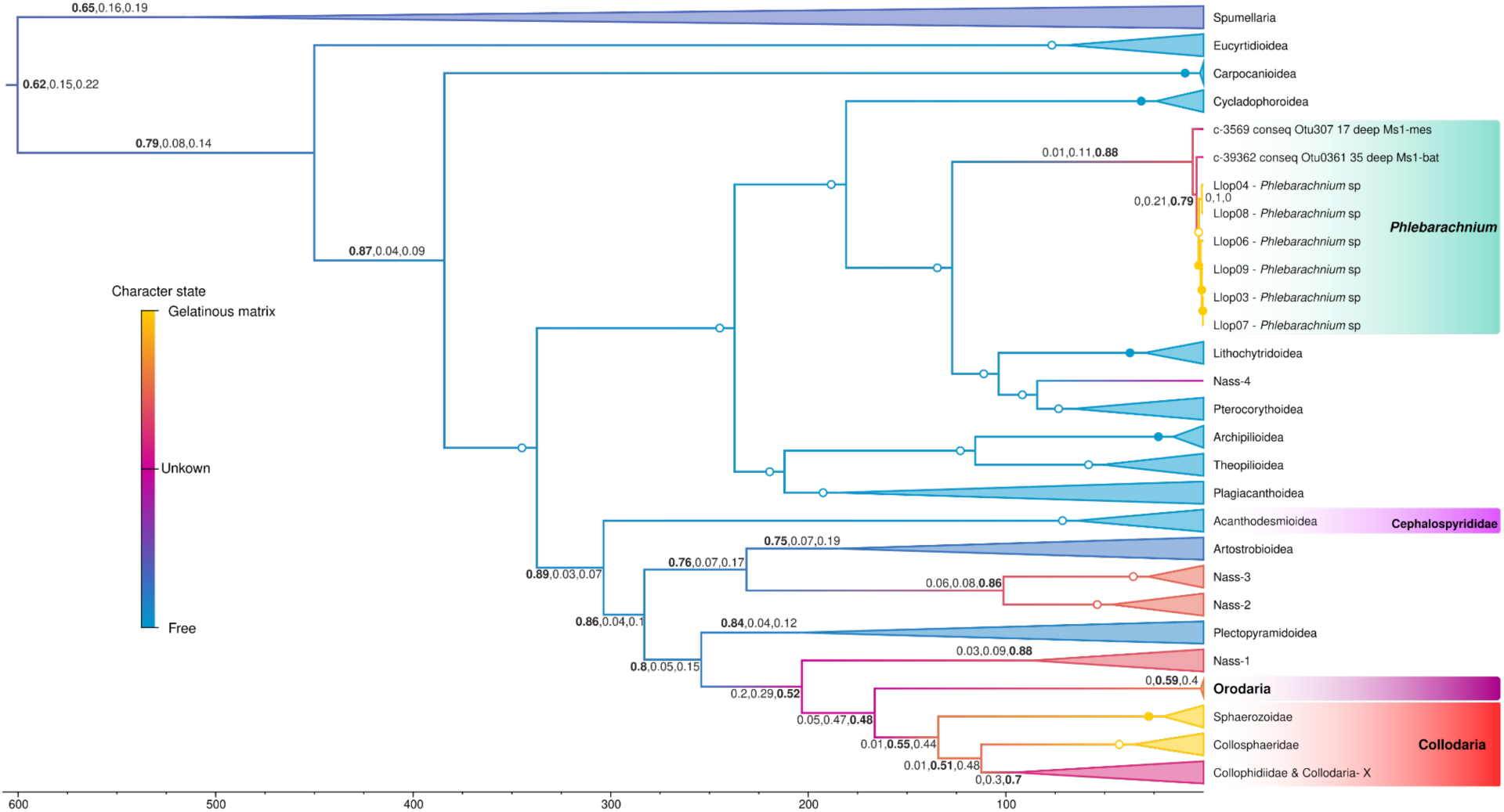
Bayesian ancestral state reconstruction performed under a birth death model implemented in BEAST2 with 10 million generations sampled every 1000 steps and a 25% burnin. Values above branches represent the probability of the absence of a gelatinous matrix ancestral state, of a gelatinous matrix ancestral state and of an unknown state (considered as a third state methodologically) respectively. When full support for either state, probabilities are represented by a full circle. Hollow circles represent posterior probabilities equal or bigger than 0.90 for the given state and smaller than 0.05 for the two other states.

**Figure S3.**
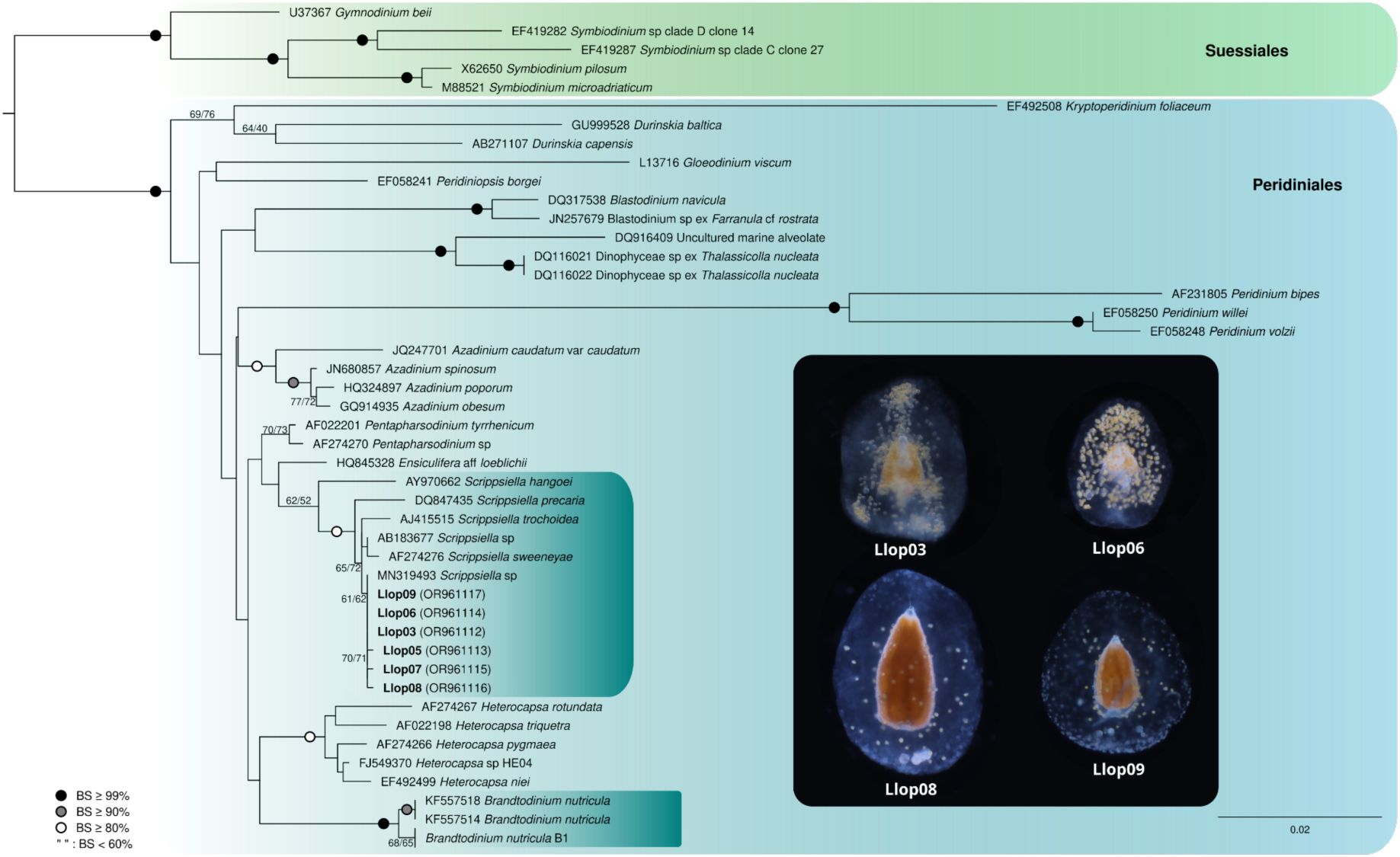
Molecular phylogeny of Perinidiales, with special focus on symbiotic dinoflagellates inferred from the 18S rDNA (71 taxa and 1803 aligned positions) using the sequences published in [21] as reference. Values at nodes represent RAxML-ng (GTR+Gamma; tree shown in figure), IQ-Tree (GTR+F+F4; 10 runs) bootstraps values (BS; 1000 bootstrap each). Only values aboce 60 BS are shown. Hollow circles at nodes indicate BS ≥ 80, filled grey circles BS ≥ 90 and filled black circles BS ≥ 90 and filled black circles BS ≥ 99.

**Figure S4.**
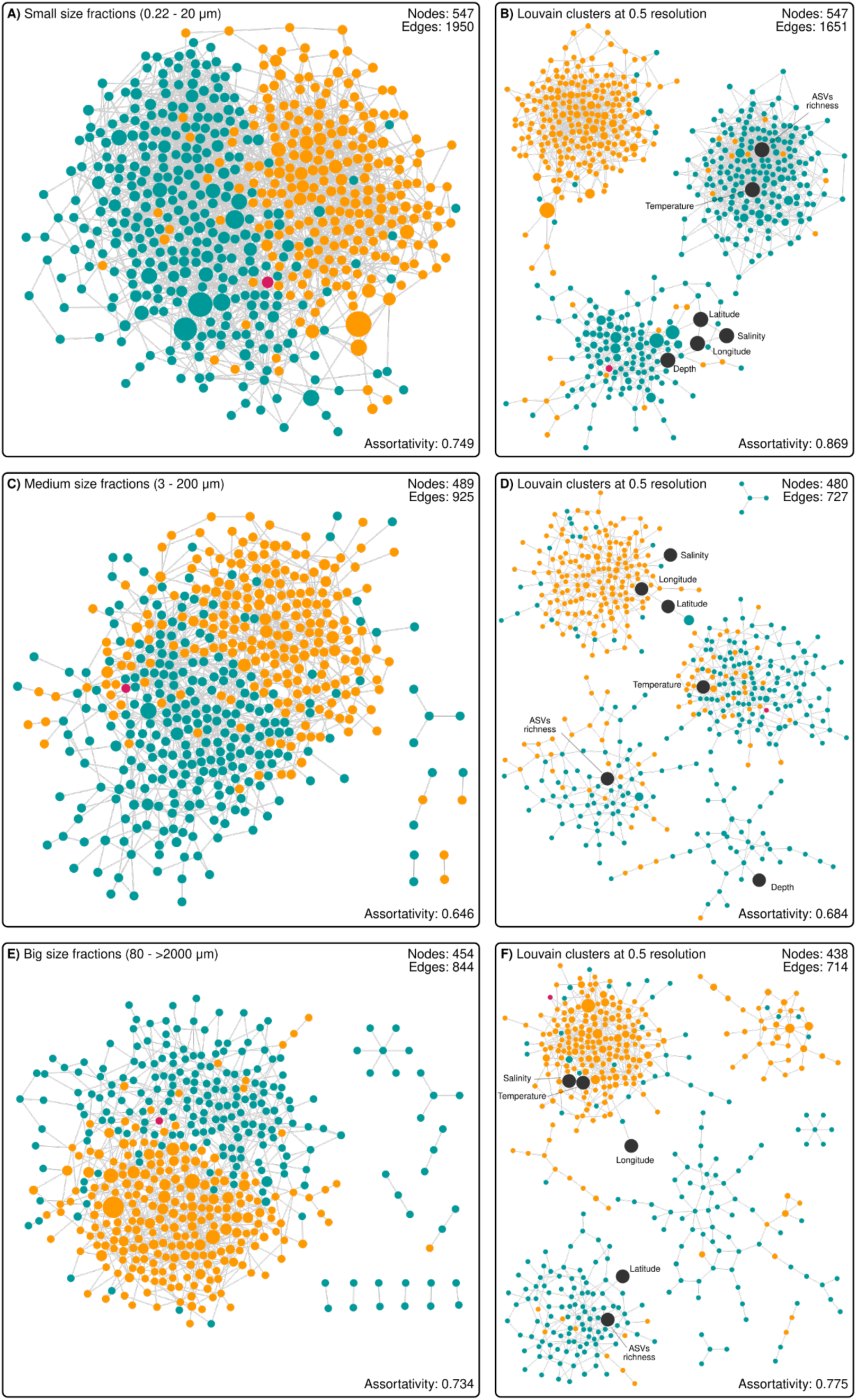
Detailed size-fraction co-occurrence networks of Radiolaria across the global oceans highlighting gelatinous and non-gelatinous amplicons (nodes) extracted from the EukBank dataset shown in Fig. 6. Left panel (A, C, D) shows the different arbitrary size-fractions and corresponding louvain clusters on the right panel (B, D, E, respectively). For further details, see Fig. 6 legend.

**Figure S5.**
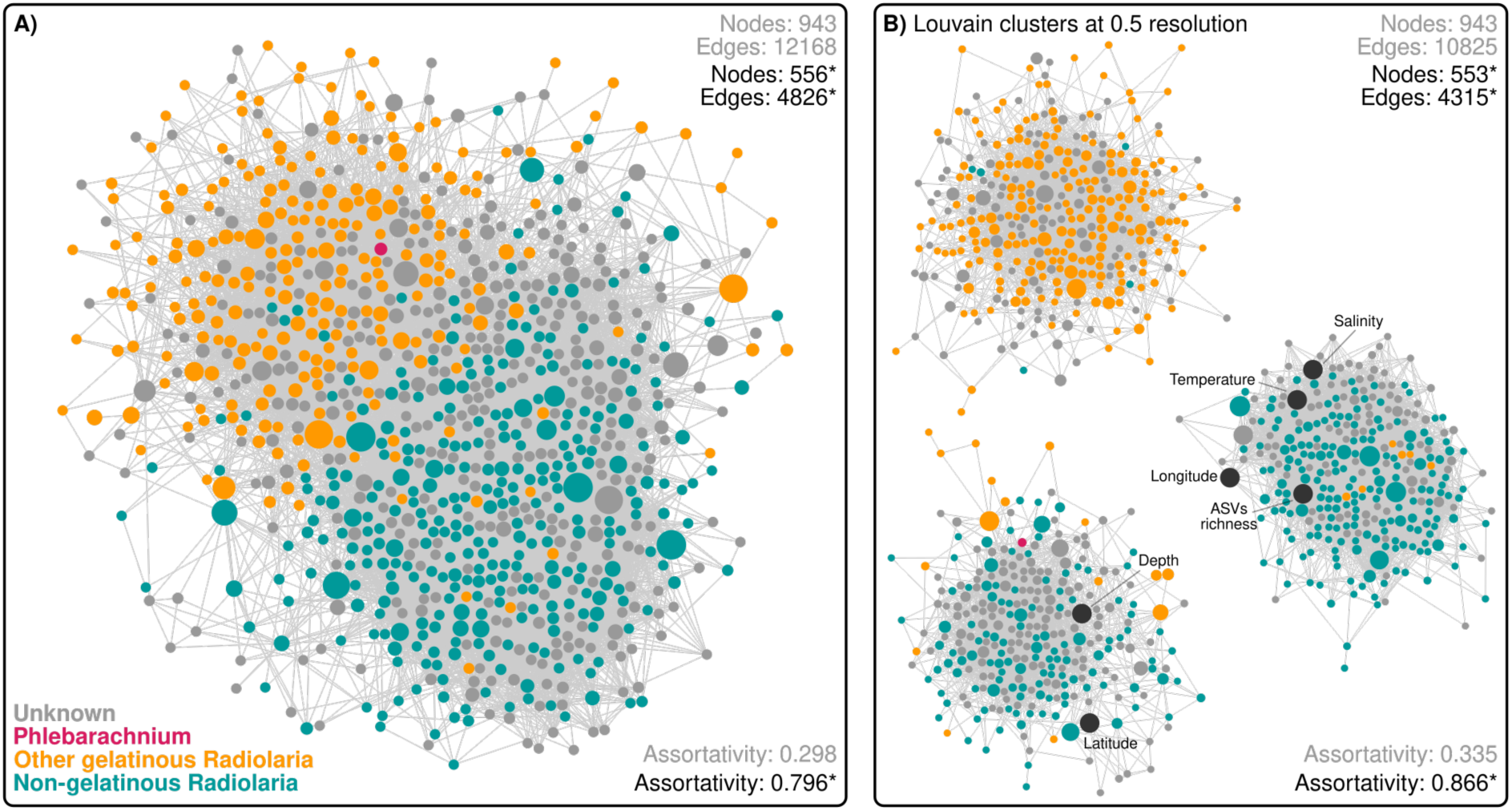
**A**) Co-occurrence network of Radiolaria across the global oceans highlighting gelatinous and non-gelatinous amplicons (nodes) extracted from the EukBank dataset and including amplicons whose morphology remains unknown. **B**) Louvain clusters of the original network shown in A) explaining community structure and co-occurrence. Given that assortativity is property of nodes with given attributes, the absence of attribute is interpreted as a third attribute, and thus assortativities are given for the shown graph (in gray) and for the same graph after removing nodes with unknown attributes (in black, with an asterisk).

